# Global maps of lake surface water temperatures reveal pitfalls of air-for-water substitutions in ecological prediction

**DOI:** 10.1101/2022.08.22.504769

**Authors:** David W. Armitage

## Abstract

In modeling species distributions and population dynamics, spatially-interpolated climatic data are often used as proxies for real, on-the-ground measurements. For shallow freshwater systems, this practice may be problematic as interpolations used for surface waters are generated from terrestrial sensor networks measuring air temperatures. Using these may therefore bias statistical estimates of species’ environmental tolerances or population projections – particularly among pleustonic and epilimnetic organisms. Using a global database of millions of daily satellite-derived lake surface water temperatures (LSWT), I trained machine learning models to correct for the correspondence between air and LSWT as a function of atmospheric and topographic predictors, resulting in the creation of monthly high-resolution global maps of air-LSWT offsets, corresponding uncertainty measures, and derived LSWT-based bioclimatic layers for use by the scientific community. I then compared the performance of these LSWT layers and air temperature-based layers in population dynamic and ecological niche models (ENM). While generally high, the correspondence between air temperature and LSWT was quite variable and often nonlinear depending on the spatial context. These LSWT predictions were better able to capture the modeled population dynamics and geographic distributions of two common aquatic plant species. Further, ENM models trained with LSWT predictors more accurately captured lab-measured thermal response curves. I conclude that these predicted LSWT temperatures perform better than raw air temperatures when used for population projections and environmental niche modeling, and should be used by practitioners to derive more biologically-meaningful results. These global LSWT predictions and corresponding error estimates and bioclimatic layers have been made freely available to all researchers in a permanent archive.

## Introduction

Ecologists regularly make use of climatic data for modeling species distributions, projecting population dynamics, and reconstructing prehistoric environments. Though long-term time series from individual weather stations are becoming increasingly available, their use is limited by large gaps in their spatiotemporal coverage. To address this, researchers have developed climate surfaces comprising statistically-interpolated spatial grids of climate data estimated using aggregated weather station measurements. The most popular climate surfaces are global in extent with resolutions of approximately 1 km^2^ (Fick & Hijmans, 2017; Karger *et al*., 2017). From these, annual summary statistics called *bioclimatic variables* are calculated for a grid cell. These biologically-relevant variables measure annual averages, variation, and seasonal extremes of temperature and precipitation.

Though widely used in statistical and simulation modeling, bioclimatic variables and the climate surfaces from which they are derived are still only approximations of actual on-the-ground climate conditions. In particular, aquatic ecologists working in surface waters above the thermocline often turn to land-based weather station or bioclimatic data as proxies for *in situ* temperature measurements (e.g. Lopes *et al*., 2017; Heneidy *et al*., 2019; Armitage & Jones, 2020; Zhang *et al*., 2020; Kariyawasam *et al*., 2021). While such data are commonly acknowledged to be imperfect approximations of surface water temperatures, the correspondence between air and lake surface water temperatures (LSWT) has only been measured for a small number of large lakes (Mccombie, 1959; Toffolon *et al*., 2014; Roberts *et al*., 2017; Piccolroaz *et al*., 2018), and it is unclear how generally representative the interpolated bioclimatic descriptors of these air temperatures are of LSWT. Exploring the exchangeability of *in situ* and interpolated climate measurements is important, as many pleustonic and epilimnetic organisms such as macrophytes, phytoplankton, and larval insects are of interest in species distribution modeling, population dynamic forecasting, or serve as paleoclimatic bioindicators. Yet these and other applications all make use of temperature data — most commonly in the form of *in situ*, satellite-derived, or interpolated air temperatures from points corresponding to water body surfaces.

Although sophisticated, highly-accurate models exist for estimating lake surface temperatures from air temperatures (Toffolon *et al*., 2014; Read *et al*., 2019), they tend to require parameter tuning with *in situ* surface temperature data or site-specific contextual data such as water clarity, wind conditions, and bathymetry. These requirements limit their applicability outside of well-surveyed regions. Without these data, aquatic ecologists are often left to accept the limitations of air temperature proxies and move forward with their modeling. However, new global-scale LSWT databases have recently been developed that permit a large-scale quantification of air-water temperature relationships through which one can assess the consequences of air-for-water temperature substitutions and develop statistical models to account for the potential mismatch. Such corrections may be useful for deriving more precise inference from the ecological models in which they are used.

The objectives of this study are to gather and link existing data on lake surface water temperatures and air temperatures in order to statistically assess the correspondence between *in situ* or interpolated air temperatures and water surface temperatures. I specifically investigated the correspondence between daily and monthly mean air and satellite-measure LSWT to identify the conditions under which air temperatures can serve as proxies for LSWT. I then trained machine learning models to accurately correct for this air-water temperature mismatch and use the results to derive new global monthly layers of LSWT and its associated bioclimatic variables. I conclude by examining how these air-for-water substitutions influence population dynamic forecasting, environmental niche models, and ENM-based estimates of physiological thermal responses.

## Methods

### Linked air-LSWT dataset construction

I began by gathering data from the GloboLakes lake surface water temperature database (Groom *et al*., 2014), which is a dataset of satellite-derived daily average lake surface temperatures collected as frequently as possible over the period of June 1995 to December 2016 in NetCDF format (url:http://www.laketemp.net/home_GL/). I took the spatial average of each lake’s pixels (0.025 ř resolution) at each daily observation to represent its daily mean surface temperature. Median values yielded similar results but are not presented here. Lakes with time gaps of more than 60 consecutive days were omitted from analysis. I then censored each lake’s time series before 2006, as exploratory analysis revealed inconsistent temporal data coverage prior to this year. I also removed lakes possessing fewer than 1000 data points, resulting in a total of 905 lakes in the dataset (Fig. 1). Time series from these lakes collectively spanned over 1.5 million days, with each lake having an average of 1793 daily observations (median = 1680, sd = 565) throughout the 11-year period. All analyses were carried out in the R 4.x language (R Core Team, 2021).

**Figure 1:**
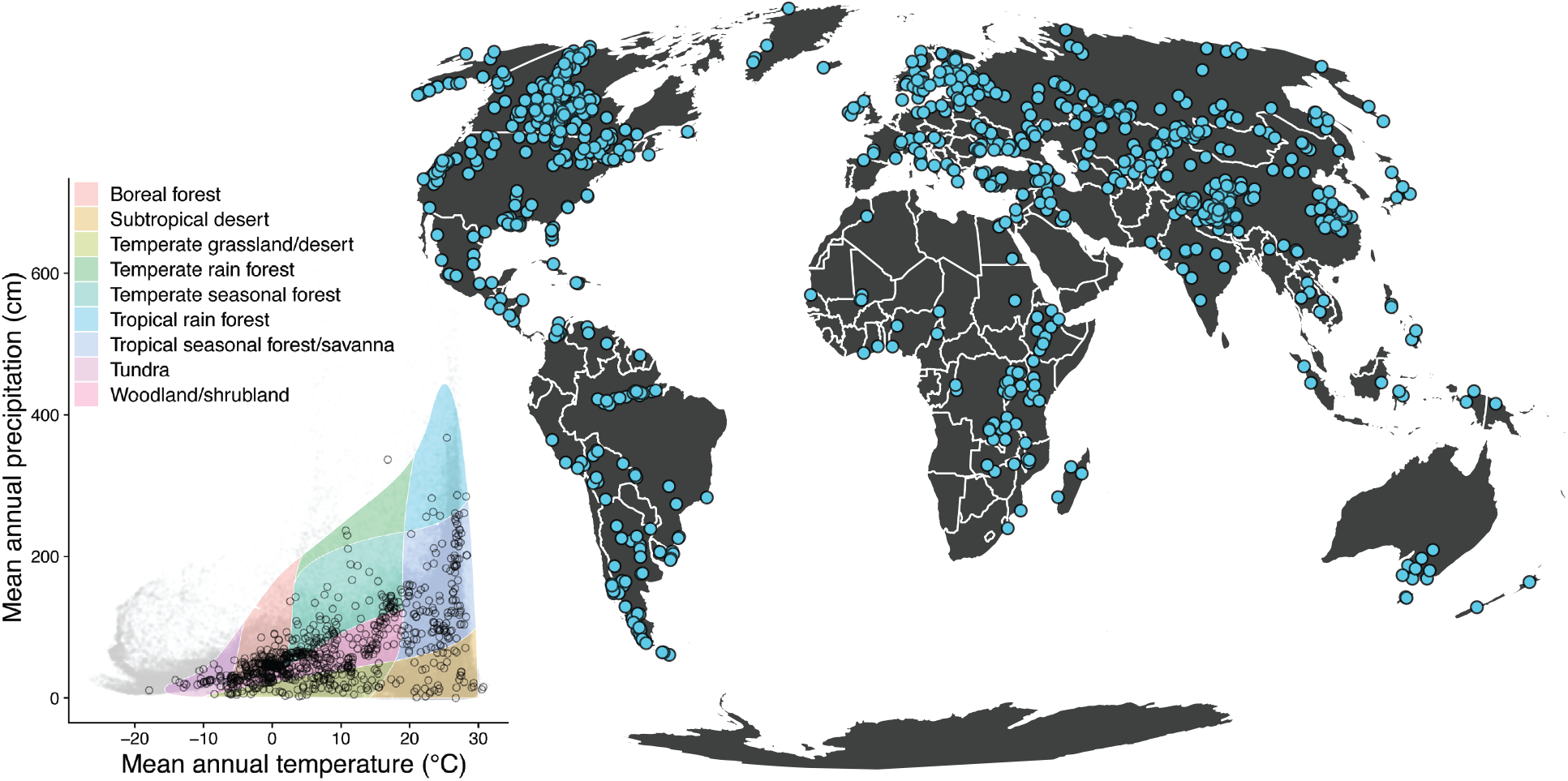
Map of lake locations obtained from the Globolakes database. Whittaker plot inset shows the distribution of lakes across (open black points) across the world’s major biomes. Light gray points show the same distribution of 100000 randomly-sampled background points.

From the coordinates of each lake (a point near its center) on a particular date, I used the R package RNCEP (Kemp *et al*., 2012) to extract a coarse-gridded daily mean 2m air temperature from the NCEP-DOE Reanalysis II data product (Kanamitsu *et al*., 2002). The spatial resolution of these air temperatures is coarse (2.5 °x 2.5 °at the equator), but was sufficient for exploring the relationship between daily air and LSWT measurements. For the same set of lakes, I also obtained fine-scale monthly-averaged data from the CHELSA (Karger *et al*., 2017; Karger & Zimmerman, 2018) climatological database (version 2.1). The spatial resolution for these data are approximately 30 arc seconds, corresponding to 1km at the Equator. For each month between January 2006 and December 2016, I extracted the following measurements from the CHELSA database: mean, maximum, and minimum daily 2m air temperatures (°C), vapor pressure deficit (Pa), near-surface wind speed (m s^-1^), precipitation amount (kg m^-2^ month^-1^/100), potential evapotranspiration (kg m^-2^ month^-1^), and near-surface relative humidity (%). Additionally, each lake’s latitude (°) and elevation (m above sea level) was obtained. For each year-month combination, I calculated maximum, minimum, and mean LSWT. Maximum and minimum temperatures were taken as the 95th and 5th percentiles of monthly lake surface water temperatures, respectively.

### Relationships between air and LSWT

With these data, I explored the relationship between measured air temperatures and lake water surface temperatures (both daily and monthly averages) using generalised additive modeling (GAM). The majority of these air-water relationships appeared flat at air temperatures below 0°C, increased approximately linearly with air temperature above freezing, and then flattened again at high air temperatures in a logistic-like fashion. To account for this nonlinearity, mixed-effects GAMs were fit with the mgcv R package (Wood, 2017) using air temperature as a fixed effect and lake ID as a random effect. I next calculated the first derivative of the GAM function with respect to air temperature, *f*′(*T*_*air*_), along with its associated 95% confidence bands using the R package gratia (Simpson, 2022). This derivative was used to identify ranges of air temperatures over which the slope of the GAM function deviated from zero, indicating the state space over which LSWT covaries with air temperature.

### Random forest prediction of LSWT

Given the nonlinear and variable association between air and LSWT, my next objective was to develop a model for predicting LSWT from air temperatures by including additional meteorological and geographic covariates that are both potentially predictive of LSWT and are also straightforward to obtain for any point on Earth. I used a machine learning approach called random forests (RF) (Breiman, 2001) to predict either daily LSWT directly, or the average offsets (i.e., difference) between min, mean, and max monthly LSWT and corresponding air temperatures. Random forests is an ensemble machine learning algorithm that that generates forests of decision trees built with both random subsets of training data as well as random subsets of covariates. The outputs of these trees are then averaged to make a prediction. I chose this algorithm rather than a more standard regression-based approach as it can model complex, nonlinear responses and interactions among covariates without assuming specifying an underlying functional form. The hyperparameters required by the RF algorithm include the number of trees being generated (here, 500), the number of covariates to split at each node, and the minimal node size per tree. For regression-type RF, it is recommended to set the latter two values to 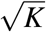 (where *K* is the number of predictor variables in the model), and 5, respectively (Wright & Ziegler, 2017). These decisions were verified by first conducting a gridded search across these hyperparameters on a subset (10%) of the data and assessing each parameter combination’s fit using 100 bootstrap resamples of the training set. To train each model, I split the LSWT dataset into 10 spatially and temporally (by year) partitioned datasets using the R package CAST (Meyer & Ludwig, 2022). Next, I measured pairwise Pearson correlation coefficients ρ between predictor variables and of those pairs with ρ *>* 0.75, only one predictor was selected at random to be retained. I used the R package ranger (Wright & Ziegler, 2017) for model fitting, which is a fast implementation of the RF algorithm for large datasets. Accuracy metrics of the fitted RF models was assessed using 10-fold cross-validation. In the case of monthly LSWT offsets, unique RF models were fit to data from each month. Predictive accuracy of each final model was assessed using an *R*^2^-like metric representing derivations of observed and predicted LSWT values from the 1:1 line (Dietze, 2017) for both in-bag (i.e., all lake samples) and out-of-bag (i.e., withheld training samples) lakes. The latter metric is more useful for identifying models that can be extrapolated to new lakes. In the case of monthly data, I obtained predicted LSWTs by adding the predicted offsets of min, mean, and maximum average monthly temperatures to the observed mean air temperatures. The fit coefficient was calculated as

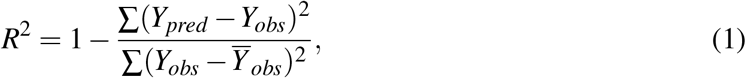

where *Y*_*pred*_ and *Y*_*obs*_ are the predicted and observed lake surface water temperatures, respectively.

### Creation of global LSWT layers

I used results from the RF models for monthly mean, min, and max temperature offsets to create global layers of predicted LSWT for each month. To do so, I assembled stacked rasters of mean monthly climatological covariates from the CHELSA database, elevation, and latitude. These objects were used to predict the per-pixel air-LSWT offset values for mean, minimum, and maximum monthly temperatures, which were subsequently used to back-calculate predicted LSWT from rasters of mean, minimum, and maximum monthly air temperatures. Spatial precision of these models was assessed following the recommendations of van den Hoogen *et al*. (2021). For each month, 100 random bootstrap samples of the training dataset were drawn and used to train RF models with their previously-selected sets of parameters and hyperparameter values. The results of these 100 models were then used to estimate spatial variance in LSWT predictions through the calculation of per-pixel standard deviations of bootstrapped model outputs. Such values are of use for practitioners when judging whether to accept the results of spatially-extrapolated model predictions.

Further, since machine learning predictions become increasingly untrustworthy as extrapolations are made beyond the predictor space of the training dataset (Meyer & Pebesma, 2022), I mapped the global coverage of spatial extrapolation outside of the dataset on a per-month basis using the *area of applicability* (AOA) metric (Meyer & Pebesma, 2021). This approach begins by centering and scaling the predictor variables of points to be predicted (here, spatial pixels on a map). Next, predictor variables are weighted based on variable importance factors (VIFs) calculated during the RF training process. This ensures that variables which are minimally important for prediction do not unduly affect the dissimilarity calculation. For each spatial pixel to be predicted, a Euclidean distance between its set of predictors and the set of training predictors is calculated, from which nearest training point in predictor space is identified. A dissimilarity index is then calculated by dividing this minimum distance by the average of all pairwise distances among the training data. The AOA is represented by those points which fall below a dissimilarity threshold, defined here as the outlier-removed maximum dissimilarity index of the training data as calculated between a training point and all remaining training points outside of its particular cross-validation fold. As AOA was calculated on a per-month basis, I also averaged these binary outcomes to generate a map showing the percentage of months a particular grid cell was within this AOA. Practitioners are encouraged to consult these mean annual maps and the corresponding individual monthly SD and AOA maps when deciding whether to use these LSWT corrections for their particular spatial and temporal requirements.

### Population dynamic modeling

Researchers may wish to use temperature-related variables to either project ecological models to locations without *in situ* data (e.g., Armitage & Jones 2020), to guide the design of experiments (e.g., Karrenberg *et al*. 2019), or — most commonly — to fit environmental niche models (ENMs) to georeferenced observations (Peterson *et al*., 2011).

I used a previously-developed population dynamic model to investigate the consequences of projecting aquatic plant population dynamics using temperature time series — either daily observed LSWT and air temperatures, or daily LSWT estimates from the random forest model.

The population dynamic model is a simplified version of a model describing the temperature and density-dependent growth of the globally-distributed floating duckweed *Lemna minor*. For details on this model, I refer readers to the original publications (Armitage & Jones, 2019). My model took the form

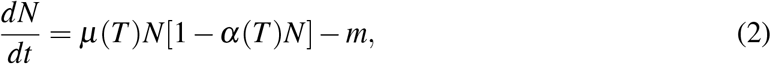

where *N* is the number of individual plants in a water body, *t* is time, µ(*T*) is the maximum temperature-dependent birth rate (per day), α(*T*) is the temperature-dependent intraspecific competition coefficient, and *m* is a constant *per capita* mortality rate. Despite its simplicity, this model has been used to predict the observed geographic distribution of the species (Armitage & Jones, 2020). For each lake in the GloboLakes dataset described above, I projected this population model with each lakes’ final ten years of lake surface water temperature measurements, as well as with the same ten years’ measurements of air temperatures and RF-predicted LSWT (from air temperature and covariates). Linear interpolation was used to fill gaps in the time series. The models were numerically solved using the R package deSolve (Soetaert *et al*., 2010). Agreement between the resulting population time series was assessed with the concordance correlation coefficient, ρ_*c*_, which measures deviation from the identity line between two time series (Lin, 1989). The coefficient, which falls between -1 and 1, is calculated as

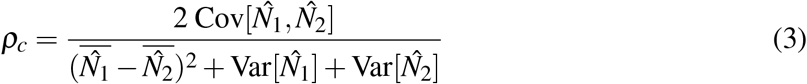

where 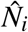 are simulated population time series generated from different temperature data (here, observed LSWT, observed air temperature, and predicted LSWT). Values of 1 indicate perfect 1:1 correspondence of two time series. Concordance correlation coefficient values were compared with nonparametric Kruskal-Wallis tests in order to assess which population time series best recaptured those based on observed lake surface temperatures.

### Bioclimatic variable calculation

I used the monthly minimum and maximum LSWT predictions and the CHELSA monthly average precipitation raster to calculate a suite of LSWT-based bioclimatic layers, which are henceforth abbreviated as *L-BIO*. These layers were calculated using the biovars function in the R package dismo (Hijmans *et al*., 2021). The correspondence between these L-BIO variables and their air temperature-based counterparts (called T-BIO) was assessed using GAM.

### Environmental niche modeling

Environmental niche models (ENMs) – also called species distribution models – make frequent use of bioclimatic variables, which are correlated against geographic observation records and used to model habitat suitability. I fit ENMs to high-density occurrence records for the aquatic duckweeds *Lemna minor* and *Spirodela polyrhiza* using sets of both L-BIO and T-BIO bioclimatic layers as predictor sets. I fit models using both the entire suite of temperature-based layers (BIO 1-11, 18, and 19) as well as a reduced set of temperature mean and variance predictors (BIO 1 and 7) for estimation of response surfaces. Details on the location records and model fitting procedure can be found in (Armitage & Jones, 2020). Points were derived from aggregated records from the Global Biodiversity Information Facility (GBIF via the rgbif package) Chamberlain *et al*. (2022) and Botanical Society of Britain and Ireland (Walker *et al*., 2010). Records span Northern Europe (including UK & Ireland) and Northern North America (USA & Canada). As the records are presence-only, I used MaxEnt (Phillips *et al*., 2006) to estimate occurrence probabilities from bioclimatic covariates.

Models were fit to spatially-thinned observations (n = 577 for *S. polyrhiza* and 909 for *L. minor*) in the ENMeval R package (Kass *et al*., 2021) using 4-fold cross-validation across various combinations of feature types (linear, quadratic, product, hinge, threshold) and regularization multipliers (2-5). Training and testing data were geographically partitioned using a nested checkerboard approach to control for spatial bias and were the same for both sets of predictors. Models fitted to T-BIO and L-BIO layers were compared using the 10% training presence omission rate (OTP10) and the continuous Boyce indices (CBI). OTP10 measures the percentage of test locations with occurrence probabilities (*p*_occ_) lower than a threshold which excludes 10% of points with the lowest occupancy probability. Here, values equal to or less than 0.1 indicate a better model fit. CBI uses a Spearman rank correlation to assess concordance between observations and predictions for ENMs using presence-only data and is therefore more appropriate for comparing MaxEnt models (Hirzel *et al*., 2006).

For each species of duckweed, I selected the best overall model formulation as that which resulted in the highest CBI score while also ensuring the OTP10 metrics were acceptably low. Response surfaces for average annual temperatures obtained from these best-fit models were compared to observed temperature-dependent growth rates measured in laboratory growth chamber assays (Armitage & Jones, 2019) by plotting the curves on a shared axis and comparing their form. For easier visual comparison, growth rate data was re-scaled to the same range as MaxEnt’s cumulative log-log output for each species. While the units of these measurements are not equivalent (growth rate versus occurrence probability), this comparison permits the assessment whether temperature-corrected data can aid ENMs in capturing the biological properties of real populations.

## Results

Both daily and monthly average LSWT-air temperature relationships were well-described by generalised additive model with a thin-plate smooth for air temperature (*T*_*air*_ edf = 9, *p* < 0.0001) and a second ridge penalty smooth for the lake random effect (*lake* edf = 0.9 (daily) and 880 (monthly), *p* < 0.0001). The overall deviance explained by this model was 81% and 96% for daily and monthly time series, respectively, indicating an acceptable fit to the data (Fig. 2).

**Figure 2:**
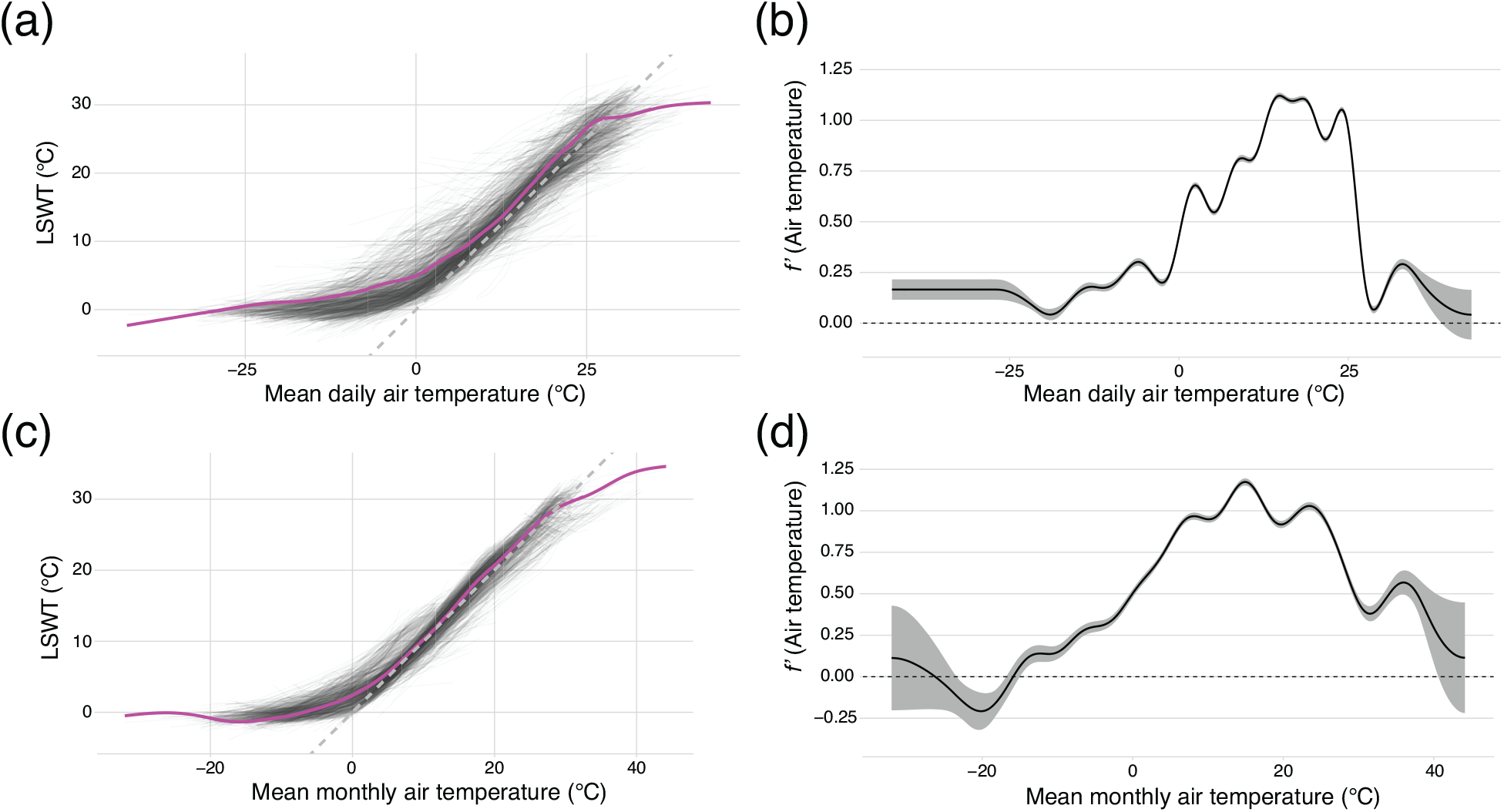
GAM fits for lakes’ **(a)** daily or **(c)** monthly air-water temperature relationships between 2006-2016. Light grey lines show GAM fits for individual lakes, while magenta line denotes the main effect of air temperature on surface water temperature. Gray dashed line shows a perfect 1:1 relationship. Panels **(b) (d)** show the corresponding the first derivatives of these GAM functions with respect to air temperature, *f*′(*T*_*air*_).

Examination of these models’ first derivatives with respect to air temperatures highlighted a general trend of near-zero slopes at air temperatures below 0°C, and a positive slope at temperatures above this point (Fig. 2). The average value of this derivative approached 1 between temperatures of 10 and 30 °C before declining again at higher temperatures, indicating a close correspondence between LSWT and air temperatures in temperate climates, but with a fair degree of nonlinearity at the extremes, as well as a non-trivial amount of spread away from this line throughout the temperature range.

Random forest models trained to correct for this nonlinearity and residual variation were able to directly predict daily LSWT time series with an out-of-bag (OOB) accuracy of 0.63 and in-bag 1:1 *R*^2^ of 0.89 (Fig. S1). These values measure expected predictive ability outside of and within the training set of lakes, respectively. Variable importance metrics ranked elevation as the most important variable in the daily LSWT RF model, and the additional covariates were approximately equally ranked but nonetheless helpful for accurate LSWT prediction. RF models for monthly LSWT-air temperature offsets had higher accuracy, with a mean OOB accuracy of 0.86 (range = 0.78-0.93) for mean temperatures, 0.80 (range = 0.72-0.87) for maximum temperatures, and 0.76 (range = 0.63-0.89) for minimum temperatures (Figs S1-S4). In-bag 1:1 *R*^2^ estimates were quite high, with that of the combined monthly mean offsets equaling 0.99, and those of minimum and maximum monthly offsets at 0.99 and 0.98, respectively. Variable importance values revealed elevation and potential evapotranspiration to be the most important predictors overall, but their relative importance varied from month to month (Fig. S5).

As the OOB accuracy of the monthly RF models was satisfactory, I used them to generate predictions of global LSWT-air temperature offsets. Average annual mean temperature offsets peaked at high latitudes and high elevations, indicating that LSWT tends to be higher, on average, than corresponding air temperatures in these regions. Conversely, LSWT in the lowland tropics and deserts tended to have LSWT lower than corresponding air temperatures (Fig. 3a). This trend held in individual months’ data, as well as for maximum and minimum temperatures (Figs.S6-S8). Area-of-applicability analysis revealed a large amount of global terrestrial space which fell outside of the range of the scaled environmental predictors. In many cases, the AOA varied month-by-month, but tended to regularly exclude large regions of Saharan Africa, Western Australia, Central America, Western S. America, Western Canada, and portions of SE Asia (Fig. 3b, S9). With regard to predictive accuracy, the standard deviation of mean annual RF models trained on 100 bootstrapped datasets revealed that the standard deviation of model predictions increased toward the North Pole, peaking with a SD of 1.2 in Siberia and Northern Canada (Figs 3c, S10).

**Figure 3:**
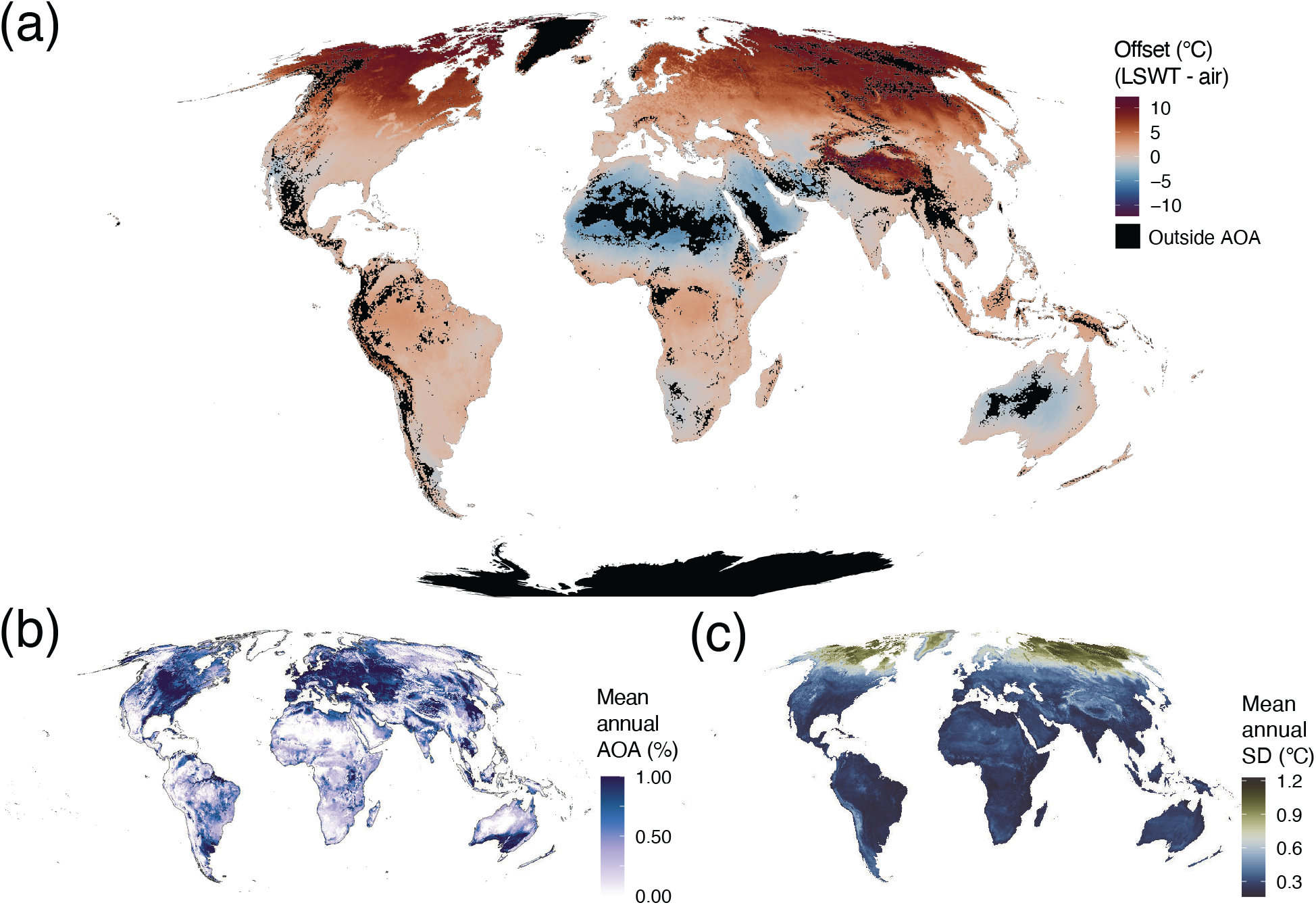
**(a)** Mean annual predicted air-LSWT temperature offsets. Values above zero indicate regions where LSWT is, on average, higher than air temperatures while values below zero indicate LSWT are lower than average air temperatures. Black regions are areas that fall outside of the area of applicability for all 12 months. **(b)** Annual mean area-of-applicability values. Shading corresponds to percentage of months a pixel falls within the AOA. **(c)** Standard deviation of bootstrapped model predictions for mean annual temperature offsets. High SD indicates model predictions are less stable and potentially less accurate.

Using different temperature time series (air, LSWT, and RF-predicted LSWT) to project the population dynamics of *Lemna minor* across the collection of lakes highlights the consequences of air-for-water temperature substitutions for population modeling. In most lakes, population projections made using observed air temperatures resulted in moderate to low correspondence with projections using observed LSWT, whereas populations projected with RF-predicted LSWT had nearly 1:1 correspondence to those made with observed LSWT (Fig. 4). A Kruskal-Wallis test of the associated ρ_*c*_ values for each pairwise comparison type found them to be significantly different (χ^2^ = 3452.3, *p <* 0.0001), with strong support for the highest concordance between observed and predicted LSWT population projections. Concordance correlation coefficients between population time series peaked at low elevations and intermediate latitudes (40-60°N/S), but those comparing predicted and observed LSWT were generally much higher over all latitudinal and elevational ranges. In sum, this evidence suggests that correcting for air-water mismatch can lead to more accurate population forecasts for surface water-dwelling organisms.

**Figure 4:**
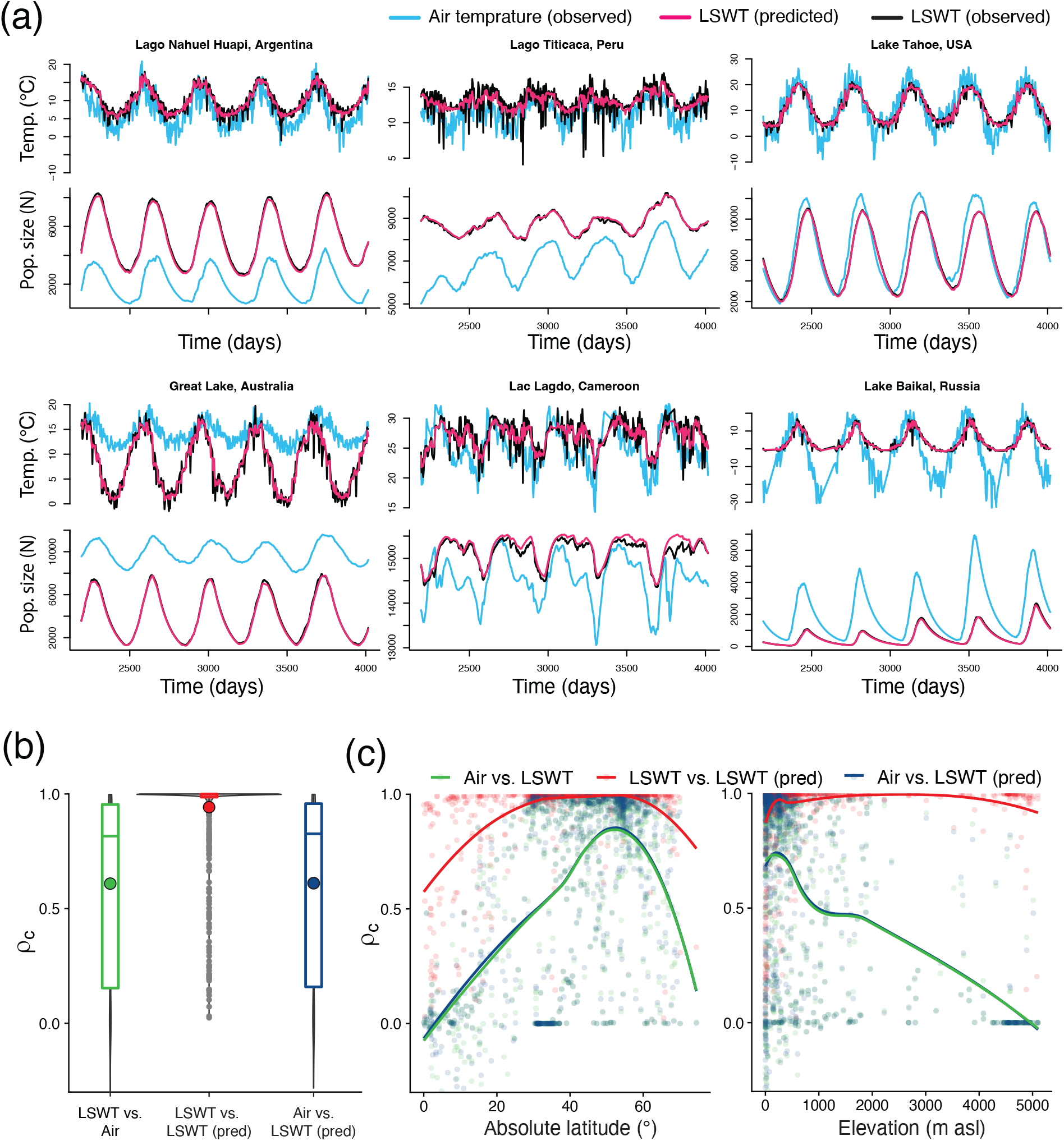
**(a)** Six example daily lake time series illustrating the correspondence between temperatures and modeled population dynamics of *L. minor*. Line colors indicate which temperature time series (observed air, observed LSWT, or predicted LSWT) were used to project populations. Example lakes were selected to span a wide latitudinal gradient. **(b)** Distribution of the concordance correlation coefficients (ρ_*c*_) comparing population dynamic projections calculated with different temperature time series for each lake in the dataset. Values of 1 indicate a perfect 1:1 correspondence of the two series. Large points denote mean ρ_*c*_ values for each comparison. **(c)** Relationship between ρ_*c*_ and latitude/elevation. Colors represent different pairwise comparisons used to calculate population time series concordance. Solid lines show GAM smooths for each comparison type.

Bioclimatic layers calculated with LSWT (L-BIO) tended to deviate from standard air-based bioclimatic layers (T-BIO) such that niche models fit for aquatic organisms using air temperatures could be unreliable (Fig. S6). To investigate this, I used MaxEnt to fit ENMs for the duckweed species *Spirodela polyrhiza* and *Lemna minor*. For almost all model specifications, fit metrics (Boyce index, 10% training presence omission rates) indicated that models trained to the L-BIO predictor set resulted in better models of habitat suitability (Fig. S12). The best-fit models for each species revealed strikingly different habitat suitability predictions for T-BIO and L-BIO predictor sets (Fig. 5). These maps, when made binary using a standard 10% omission rate threshold, resulted in markedly different range predictions between the predictor sets, with the L-BIO-trained models resulting in a relatively more accurate prediction of both species’ poleward range limits in both the US and EU (Fig. S13). Examination of the response surfaces of these best-fit ENMs for both species reveals a qualitatively closer correspondence between lab-measured thermal growth curves and ENMs fitted with L-BIO predictors than between lab-measured curves and ENMs fitted with T-BIO predictors (Fig. 5e & j). This suggests that environmental response surfaces from correlative ENMs which use lake-temperature corrected predictors more accurately captured key aspects of the duckweeds’ intrinsic thermal growth responses, particularly the species’ lower thermal limits, which appeared underestimated by T-BIO predictors.

**Figure 5:**
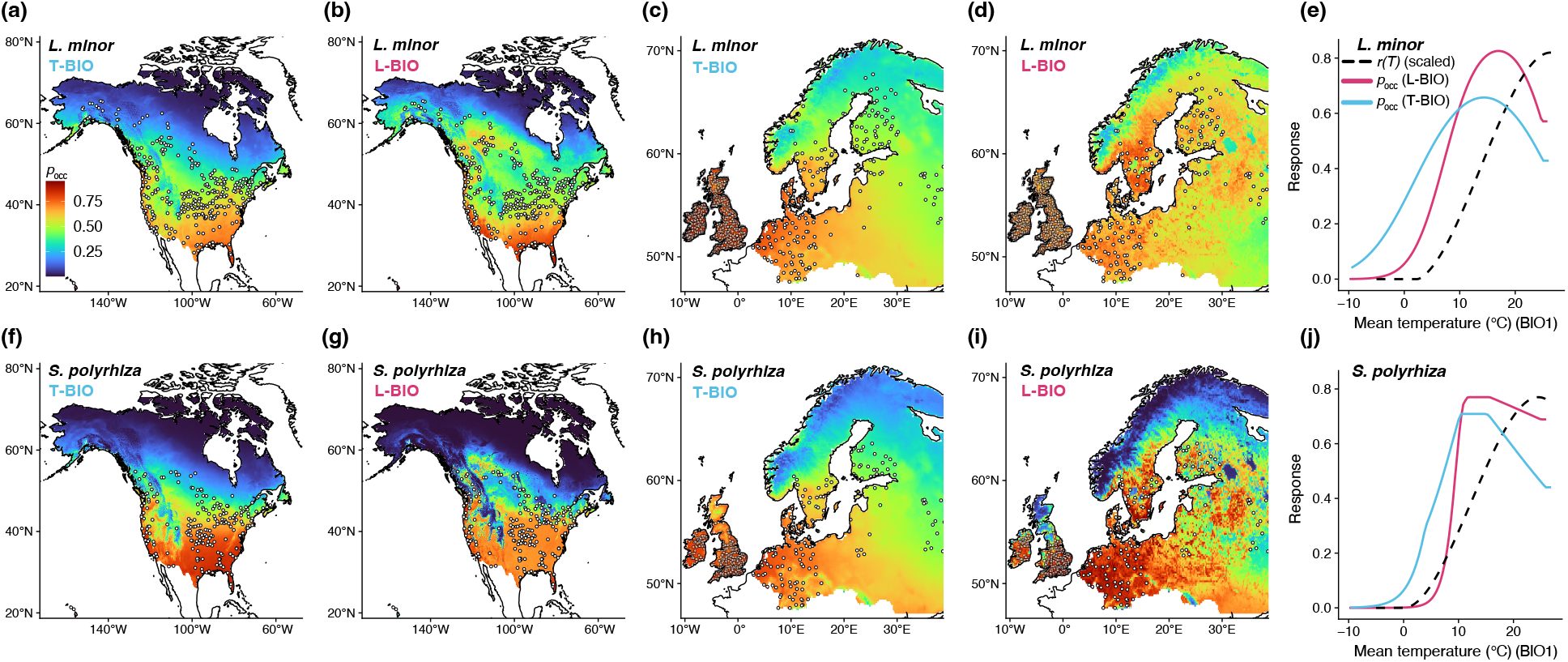
Comparisons between ENM predictions trained with air temperature-based (T-BIO) or LSWT-based (L-BIO) bioclimatic predictors for the duckweed species **(a-d)** *Spirodela polyrhiza* and **(f-i)** *Lemna minor*. Colors show cumulative log-log occurrence probabilities (*p*_occ_). Panels **(e)** and **(j)** compare the ENM-derived occurrence probabilities for average annual temperatures (BIO1) with the experimentally-measured growth curves (*r*(*T*)) for each species. Note the overall closer correspondence in form between *r*(*T*) and L-BIO responses than with the air-based T-BIO predictors.

## Discussion

While lake water surface temperatures are closely correlated with air temperatures, they do not correspond perfectly, which has motivated efforts to develop models to convert between air and LSWT. Process-based models, such as the *Air2Water* model (Piccolroaz *et al*., 2013), use mechanistic heat exchange equations between air and water to predict LSWT. While this model yields highly accurate lake surface water temperature predictions (*R*^2^ > 0.95), its use is limited to lakes from which surface temperature time series have already been collected, as these data are used in parameter tuning. Similarly, machine learning tools can perform accurate air to water conversions, again provided that a training set of water temperatures is available (Heddam *et al*., 2020). Other mechanistic models have used global hydrological models and energy balance equations to successfully predict monthly temperatures of large rivers and lakes at resolutions as high as 10 km^2^ at global scales (van Beek *et al*., 2012; Wanders *et al*., 2019). While such models are currently at the cutting edge of global river and lake temperature prediction, their utility for predicting the LSWT of smaller lakes and ponds is not clear, as they require solving computationally intensive hydrological models of regional drainage networks. Furthermore, outputs from these models currently lack bioclimatic products commonly used for environmental niche modeling.

At the other end of the complexity spectrum are simple linear corrections for water temperature. Previous work in Swiss lakes has demonstrated how correcting for elevation significantly improved air-water temperature correspondence (Livingstone & Lotter, 1998; Livingstone *et al*., 2005). Intermediate to these extremes are models that include additional covariates such as depth, volume, and surface area (Sharma *et al*., 2008; Prats & Danis, 2019), or models that capture nonlinearities at low and high temperatures (Mohseni *et al*., 1998). While such models perform very well, they also tend to require bathymetric data and local air temperature time series that are not readily available for many lakes. Thus, the approach taken here, while perhaps less precise than process-based LSWT prediction, is flexible enough to correct for lake temperatures given only the local air temperature, elevation, latitude, and some readily-available climatic predictors and can be used to model surface temperatures in lakes without pre-existing LSWT time series and bathymetric data.

The approach for LSWT estimation employed here benefits from a large global dataset and high-resolution monthly covariates to more precisely estimate correction factors for air-water conversions. While I found an overall correspondence between both daily and monthly air and lake water surface temperatures, this relationship is not 1:1, and was quite variable between individual lakes. Plots of these relationships suggest that lake surface waters in cooler climates (mean air temperatures below 5 °C) tend to be warmer than air while lakes in hot climates (*>* 25 °C) are slightly cooler than corresponding air temperatures. On the other hand, these relationships also suggest a closely-coupled relationship between LSWT and air temperatures in temperate climates. Consequently, when additional data are unavailable, *in situ* air temperatures, when measured over an entire annual cycle, can serve as a reasonable proxy for LWST, which means that studies that have employed air-for-water substitutions in temperate climates are less at risk of systematic bias owing to this mismatch. Though it is always in the practitioner’s best interest to instead correct for this air-LSWT mismatch using either their own models or the predicted air-LSWT offsets and bioclimatic layers provided herein, as I demonstrated here for populations and niche models of aquatic duckweed plants.

The ultimate goal of this exercise was to assess the consequences of replacing air temperature-based time series and bioclimatic variables with corrected versions that more closely approximated observed LSWT. When used for population modeling, predicted LSWT time series yielded population dynamics very similar to those simulated under observed LSWT. Previous studies have found that substituting water for air temperatures as state variables significantly improved predictions of vital rates and population dynamics in both animals and plants (Paaijmans *et al*., 2010; Tarkan *et al*., 2011; Sharifi *et al*., 2018). In all of these cases, however, water temperatures were directly measured using digital loggers. The RF models and bioclimatic layers provided herein can be used more broadly in situations where direct LSWT is unavailable.

Bioclimatic derivatives of the RF models’ predictions were different than those calculated with air temperatures. This again highlights the perils of using air temperatures as proxies for lake surface conditions in species distribution models. Use of L-BIO predictors in ENMs resulted in better fits of MaxEnt models to duckweed species’ occurrence records than those fit with standard bioclimatic layers. While such improvements have previously been demonstrated for models of fish distributions (Al-Chokhachy *et al*., 2013), ENMs of floating aquatic macrophytes tend to use air temperatures (Rodríguez-Merino *et al*., 2019; Armitage & Jones, 2020), which, based on these results, could miscalculate projections of climate change-driven range shifts or invasion probabilities.

Response surfaces from ENM models are often used to understand species’ environmental tolerances such as critical thermal minima and maxima. While there are significant drawbacks to such approaches (Lee-Yaw *et al*., 2022), careful comparison with ground-truthed measurements have demonstrated the vailidity of this approach in certain settings (e.g. Searcy & Shaffer, 2016; Mothes *et al*., 2019). In the case of the duckweed *Lemna minor*, experimental thermal response curves can predict its geographic distribution using air temperature-based bioclimatic predictors (Armitage & Jones, 2020). However, these predictions overestimated its poleward range margins, leading to discordance between ENM-based and lab-measured thermal responses. In contrast, comparisons between responses derived from L-BIO ENMs resulted in a much closer match to experimentally measured thermal minima for both duckweed species, which translated to improved range limit predictions (i.e., closer to the true distributional margin) than models fit to T-BIO predictors.

## Conclusion

The predictions made by environmental niche models and population dynamic models are only as good as the data from which they are derived. Owing perhaps to their ease of access, air temperatures and their bioclimatic derivatives are commonly used to model epilimnetic and pleustonic organsims. Given that air and surface water temperatures are closely coupled, this practice can still yield valuable predictions. As demonstrated here, however, such models may not be as quantitatively meaningful as those fit to the thermal conditions organisms actually experience at the water’s surface. While the modeling exercises presented here fall squarely in the domain of population ecology, I anticipate such LSWT-corrected variables to useful, for example, to the fields of paleoclimatology (Walker *et al*., 1991), harmful algal bloom prediction (Taranu *et al*., 2012), and aquaculture/fisheries management (Lehtonen, 1996). The data provided alongside this analysis include 1km-resolution monthly average, maximum, and minimum predicted lake surface water temperatures over much of the globe, the monthly predicted area of applicability of these models, and standard bioclimatic layers calculated using these predicted lake temperatures. While the addition of more lakes and years to the dataset will undoubtedly affect global LSWT offset estimates and the associated spatial error, these products can be updated by re-training the RF models described herein.

## Acknowledgements

Support for this work comes from subsidy funding to OIST. This work was conceived as a response to reviews from a previously-published manuscript, and I thank these anonymous reviewers for inspiring me to move forward on this project.

## Author Contributions

DWA conceived the ideas and designed the methodology, collected and analysed the data, and wrote the manuscript.

## Data Availability

All data products (global LSWT layers, area-of-applicability layers, and L-BIO bioclimatic layers) are available from the Dryad Digital Repository at https://doi.org/10.5061/dryad.s4mw6m990. All source code, tabular data, and fitted model objects are available on0 the Zenodo open science platform at https://doi.org/10.5281/zenodo.7014940.

