## Supplemental Figures for "Global maps of lake surface water temperatures reveal pitfalls of air-for-water substitutions in ecological prediction"

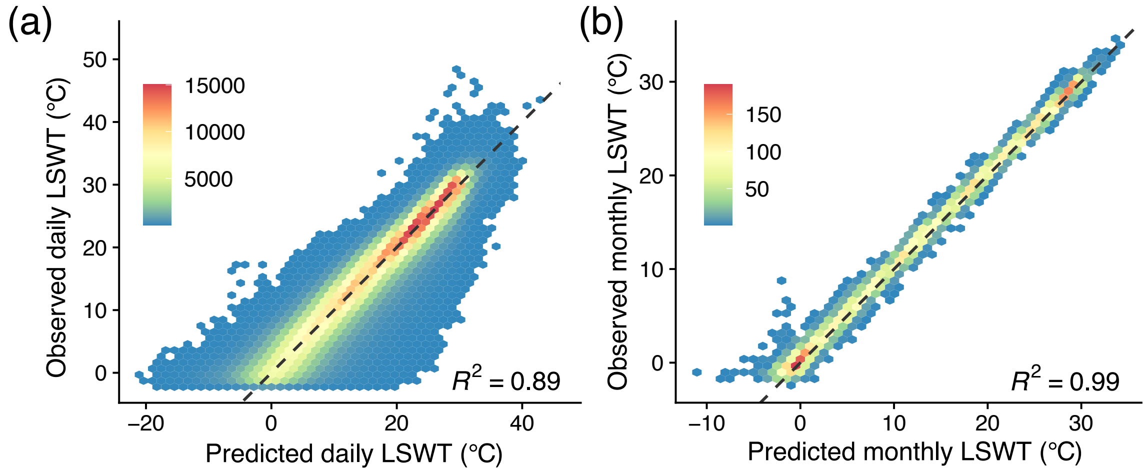


Figure S1: In-bag plots of predicted versus observed LSWT for **(a)** daily and **(b)** averaged monthly mean temperatures. $R^2$ values denote fit to dashed 1:1 line.

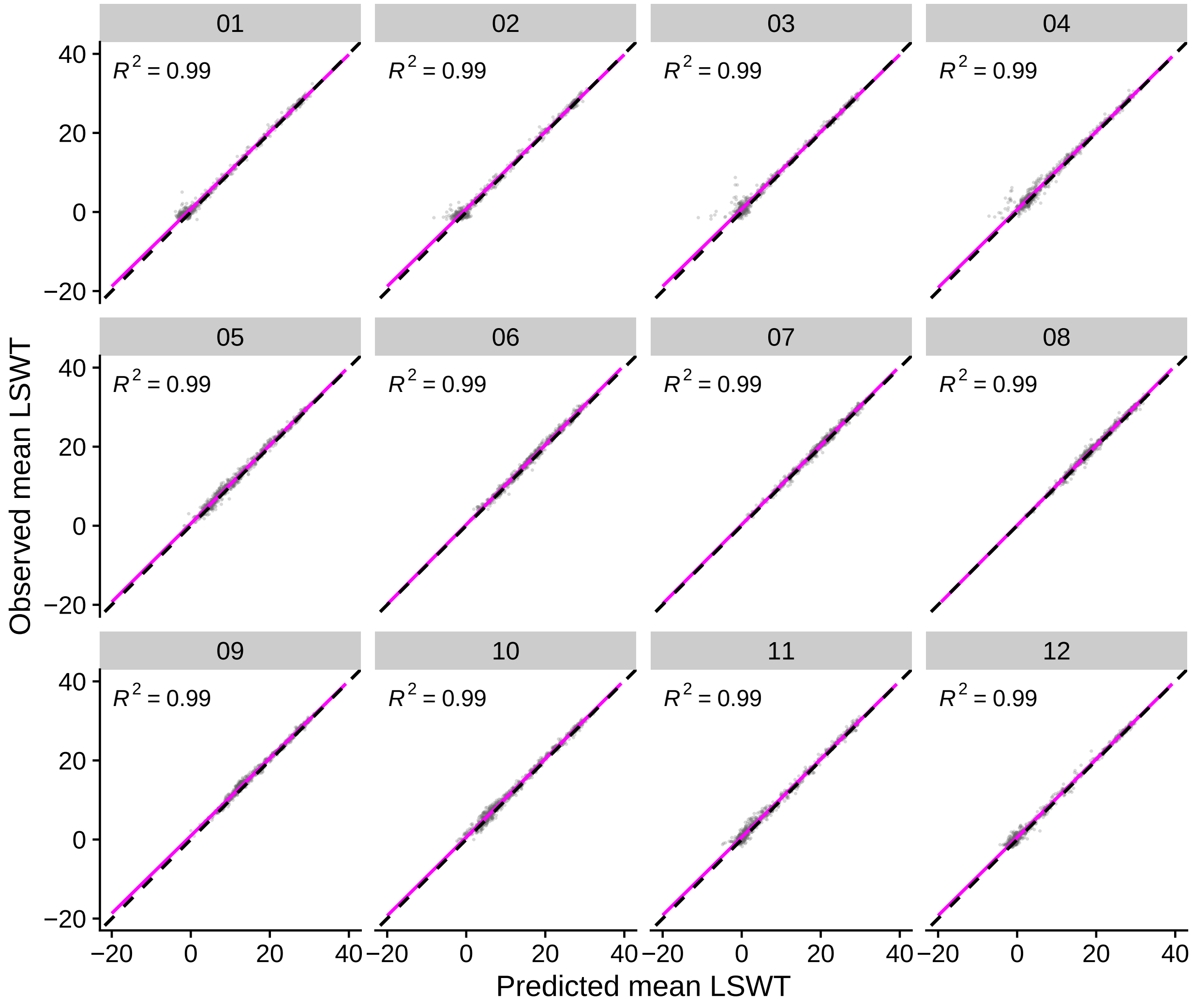


Figure S2: In-bag plots of predicted versus observed monthly mean LSWT. $R^2$ values denote fit to dashed 1:1 line. Magenta lines show linear regression fit. Facets display each numeric month of the year.

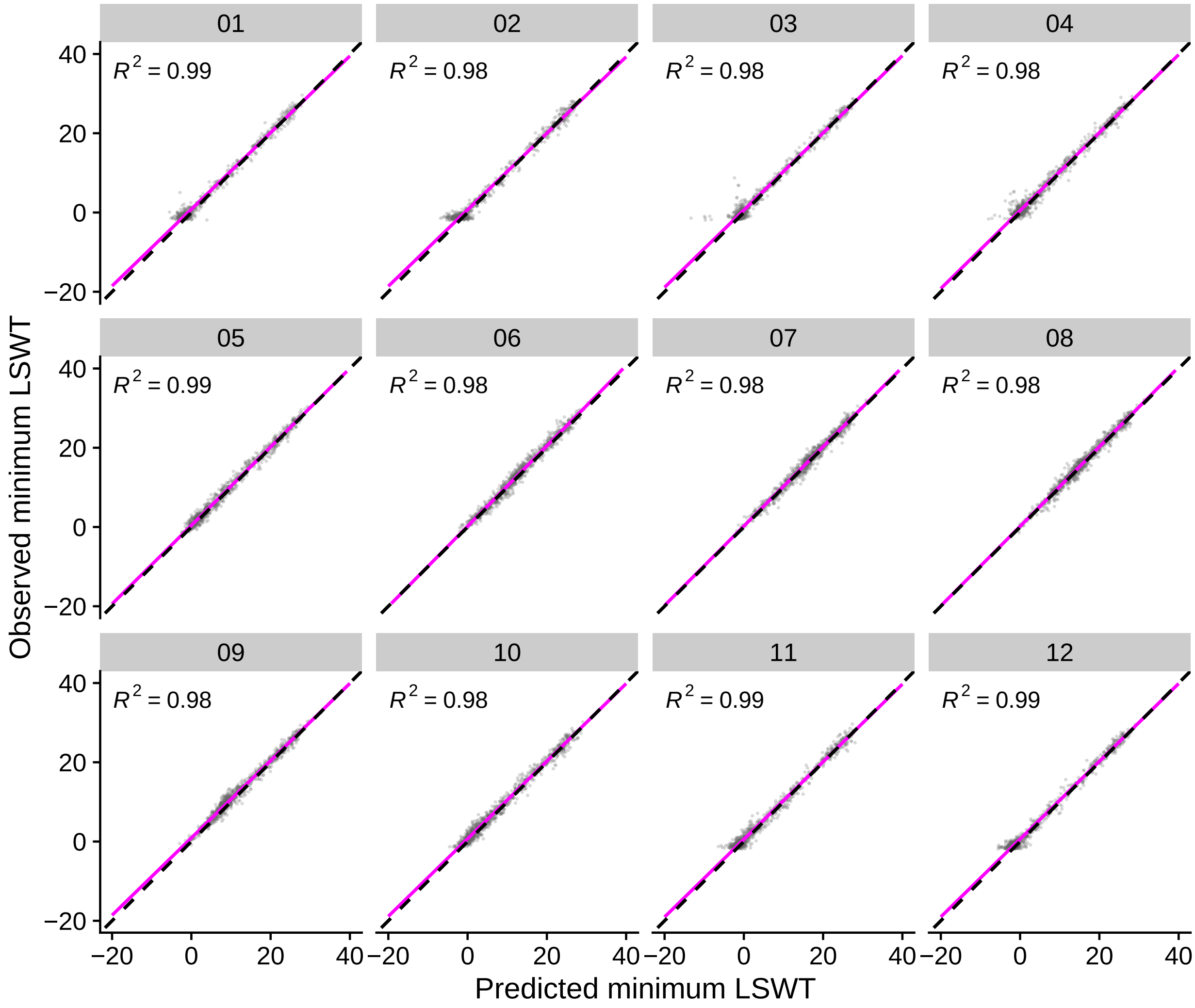


Figure S3: In-bag plots of predicted versus observed monthly minimum LSWT. $R^2$ values denote fit to dashed 1:1 line. Magenta lines show linear regression fit. Facets display each numeric month of the year.

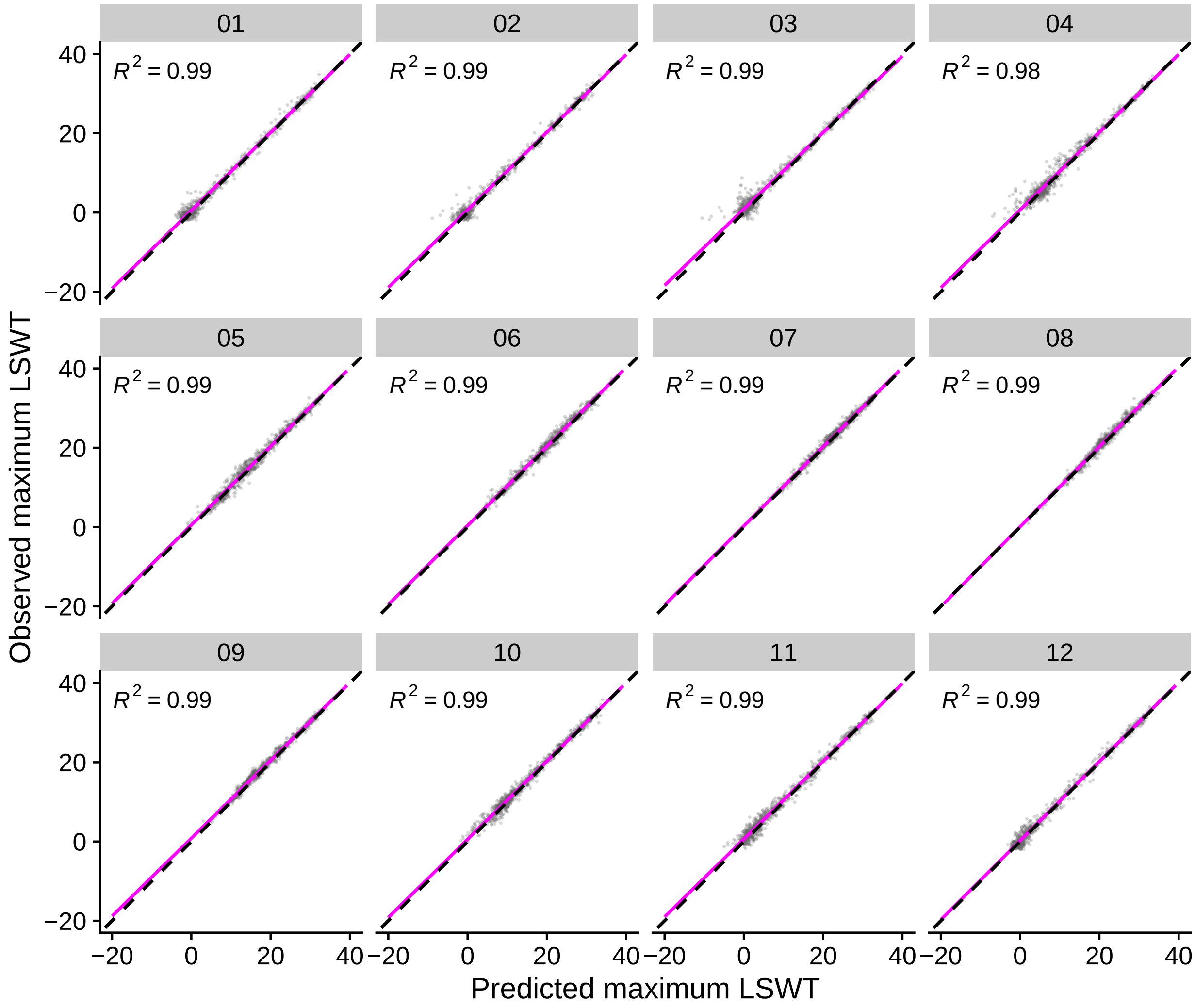


Figure S4: In-bag plots of predicted versus observed monthly maximum LSWT. $R^2$ values denote fit to dashed 1:1 line. Magenta lines show linear regression fit. Facets display each numeric month of the year.

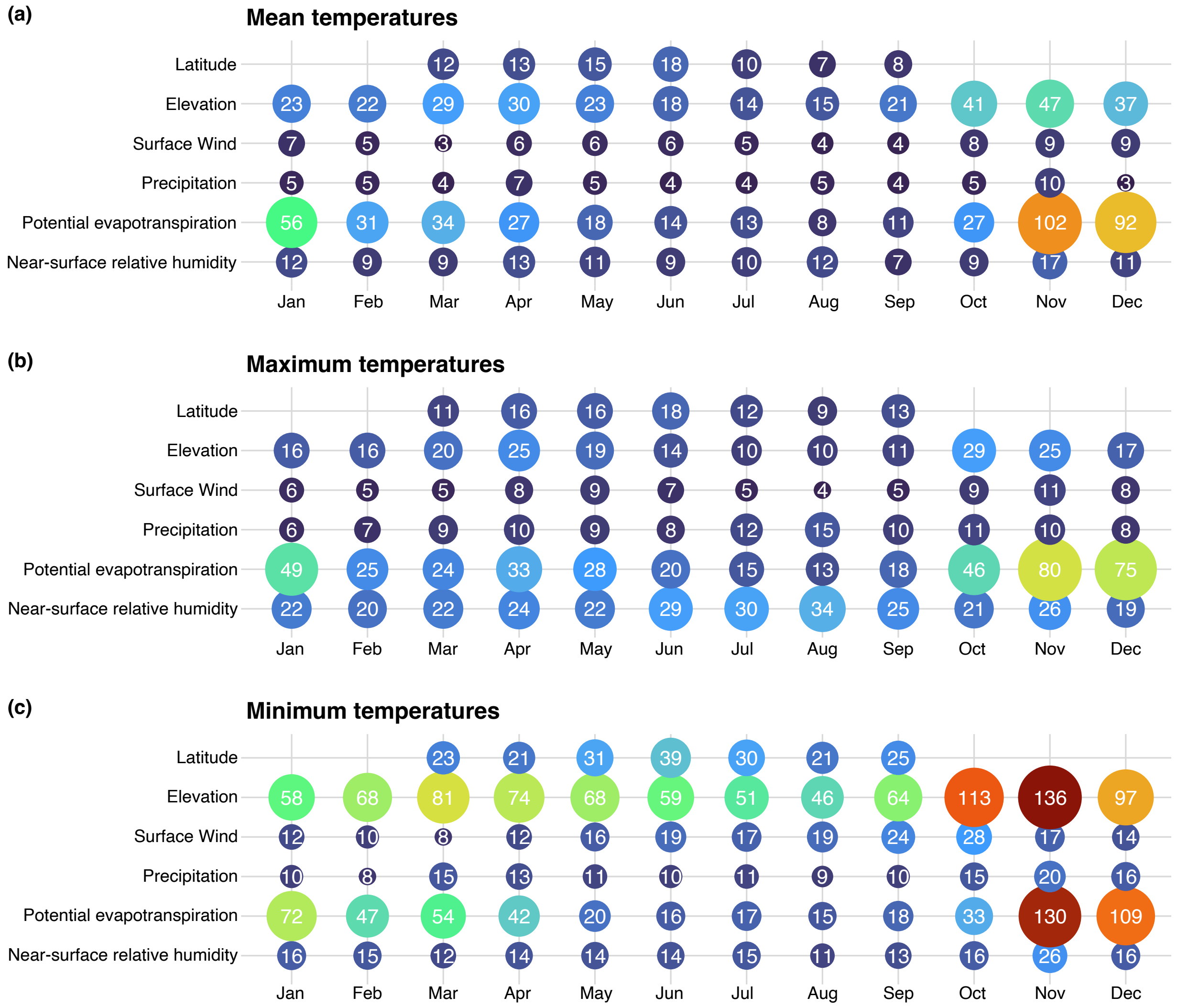


Figure S5: Variable importance plots for each monthly RF model for **(a)** monthy mean, **(b)** monthly maximum, and **(c)** monthly minimum temperature offsets. Numbers correspond to the raw variance-reduction based index for each covariate, scaled by 0.001 for convenience. Covariates with higher values are more important for predictive accuracy. Missing variables are those that were previously excluded from a particular month's RF model due to collinearity with a second, retained predictor.

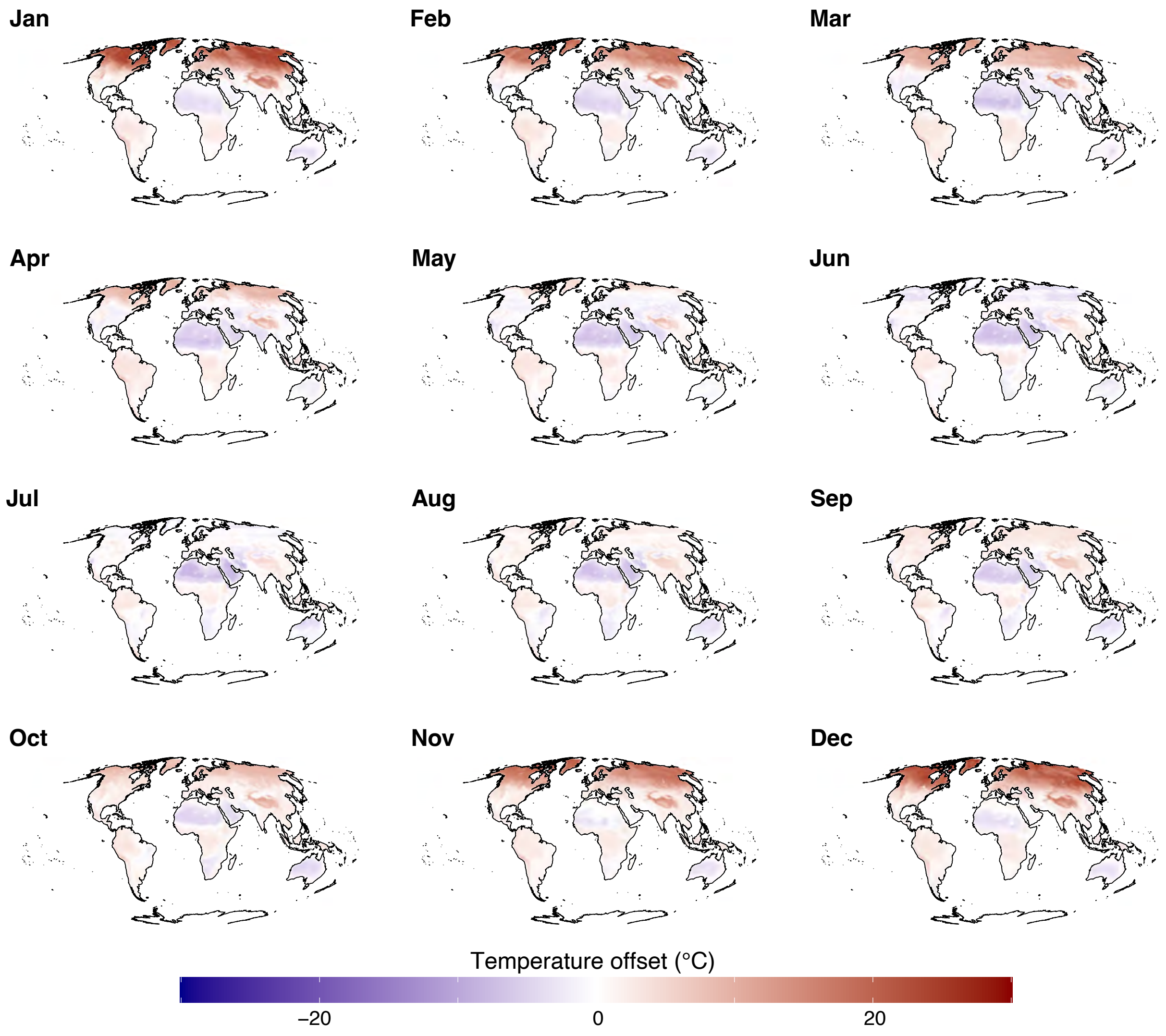


Figure S6: Global maps of RF-predicted monthly average LSWT-air temperature offsets.

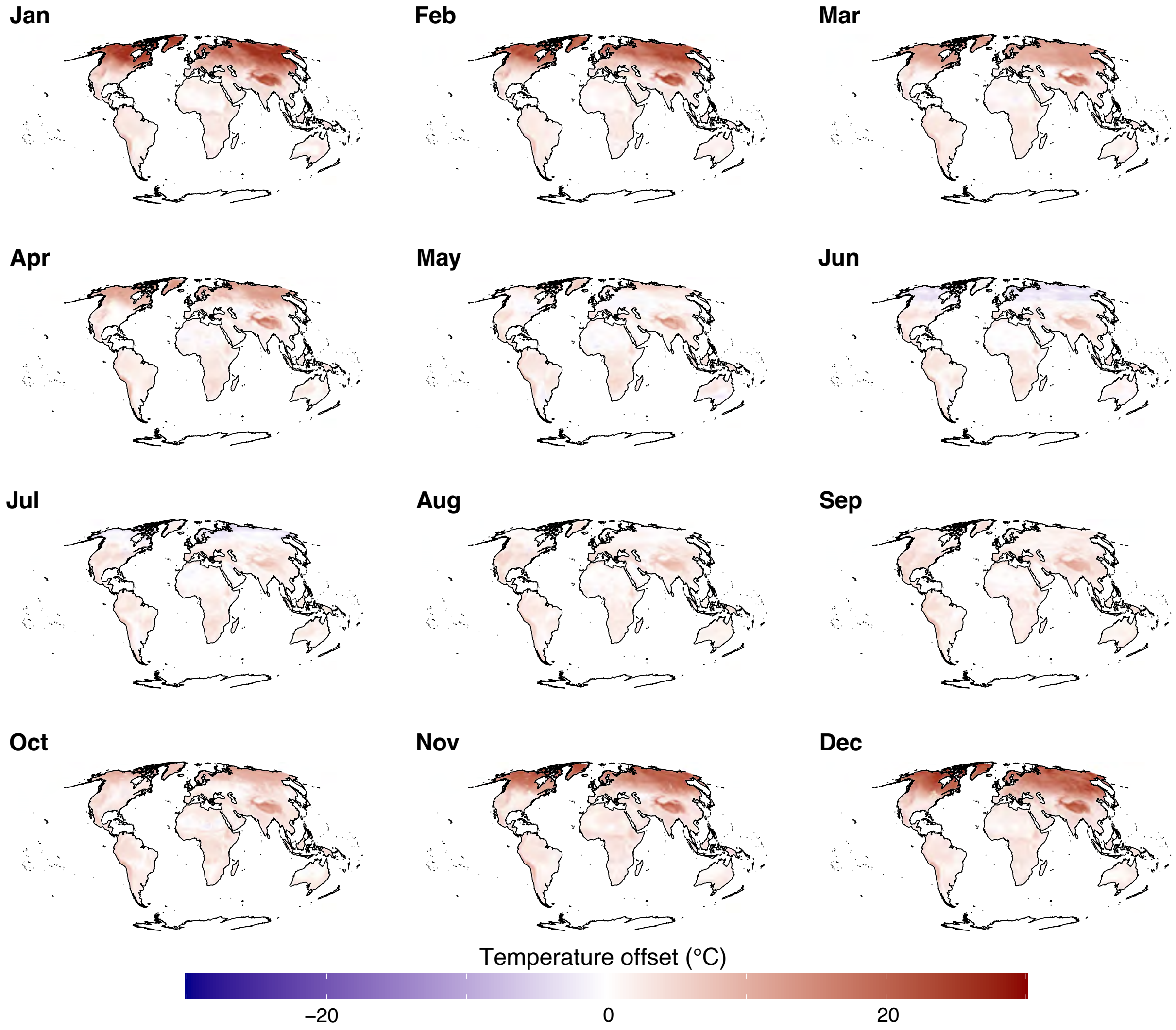


Figure S7: Global maps of RF-predicted monthly minimum LSWT-air temperature offsets.

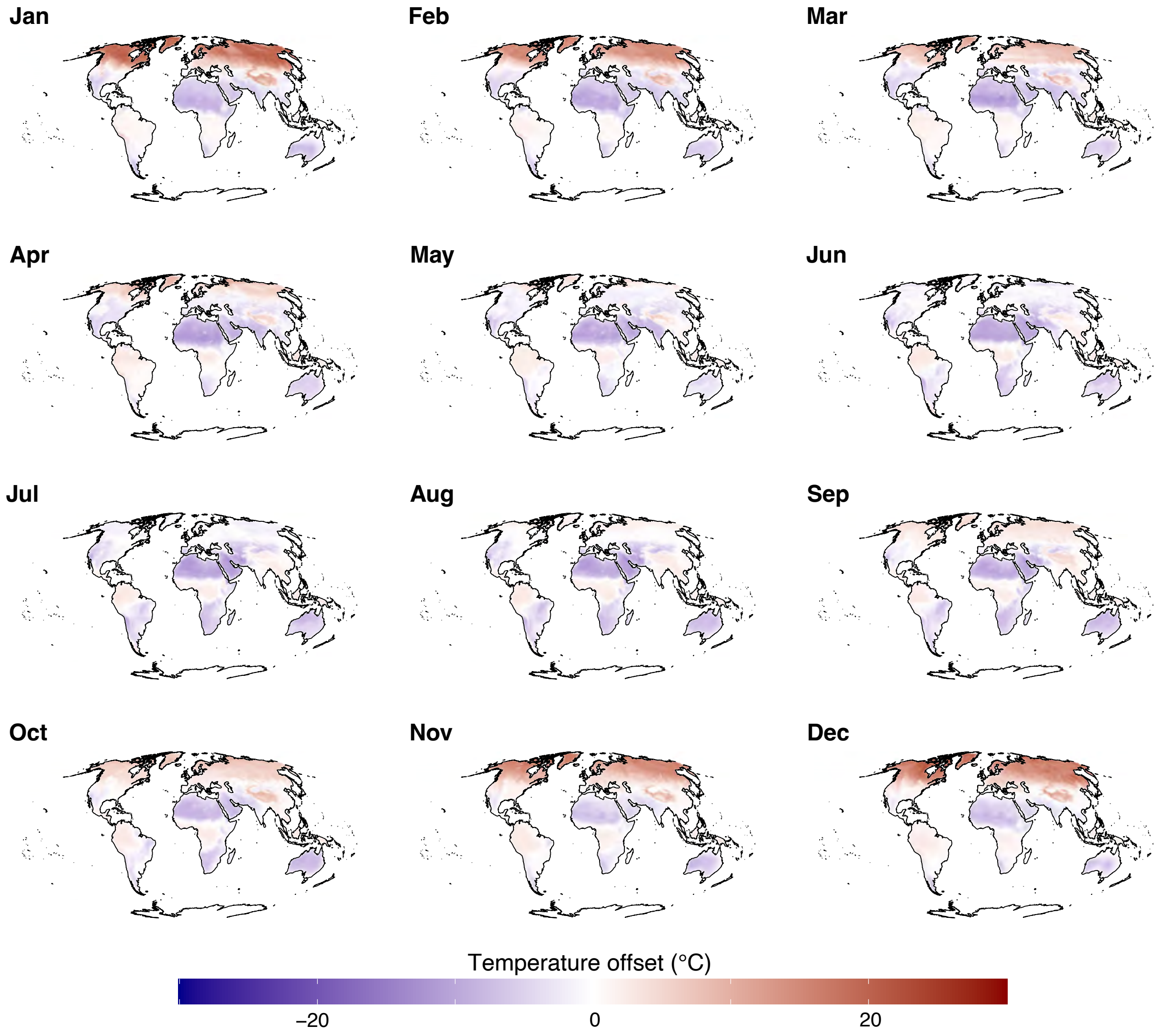


Figure S8: Global maps of RF-predicted monthly maximum LSWT-air temperature offsets.

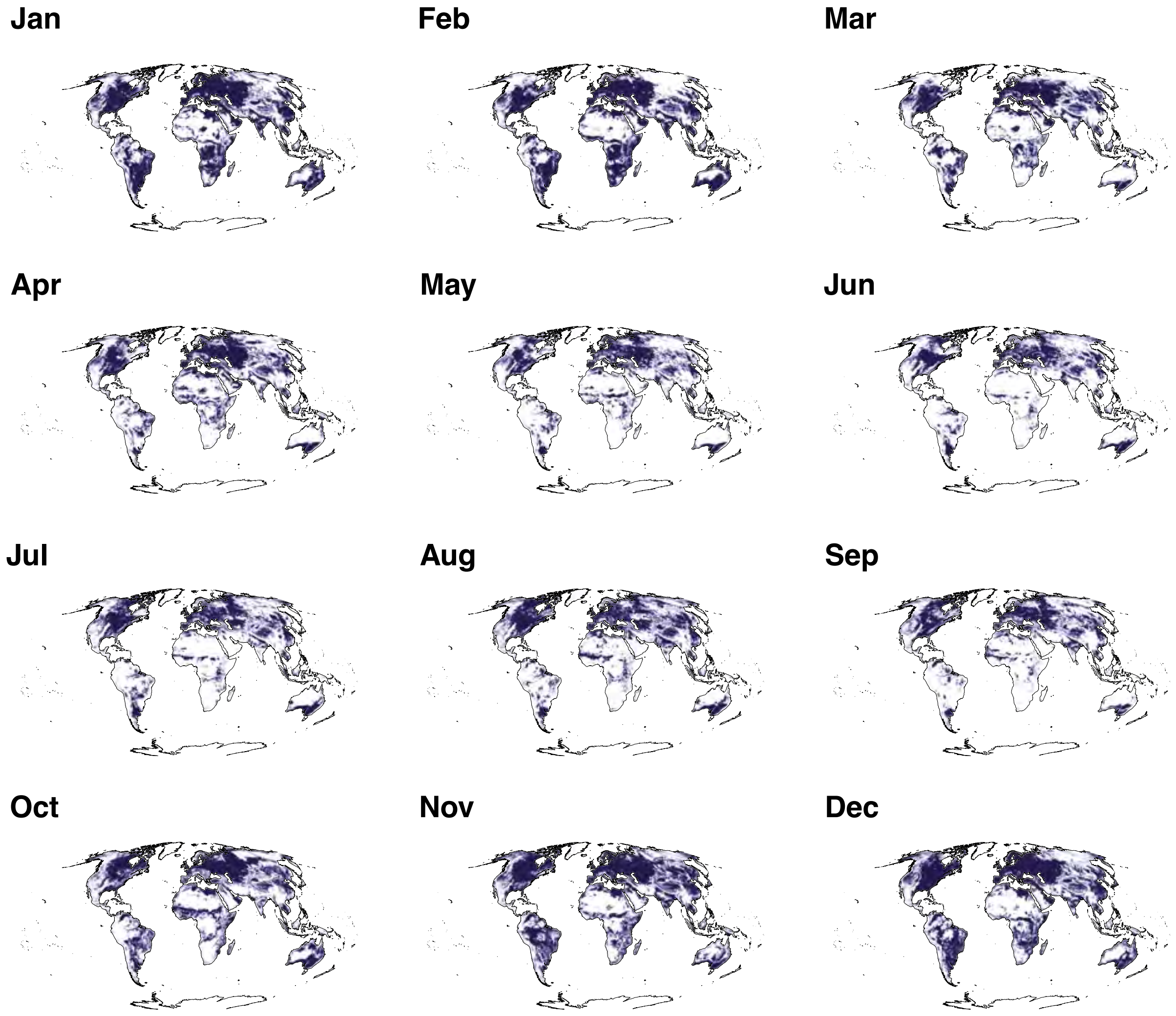


Figure S9: Global monthly maps of area of applicability (AOA). Shaded areas denote regions where the importance-weighted monthly predictor set is within the bounds of that of that month's RF model's training predictor set.

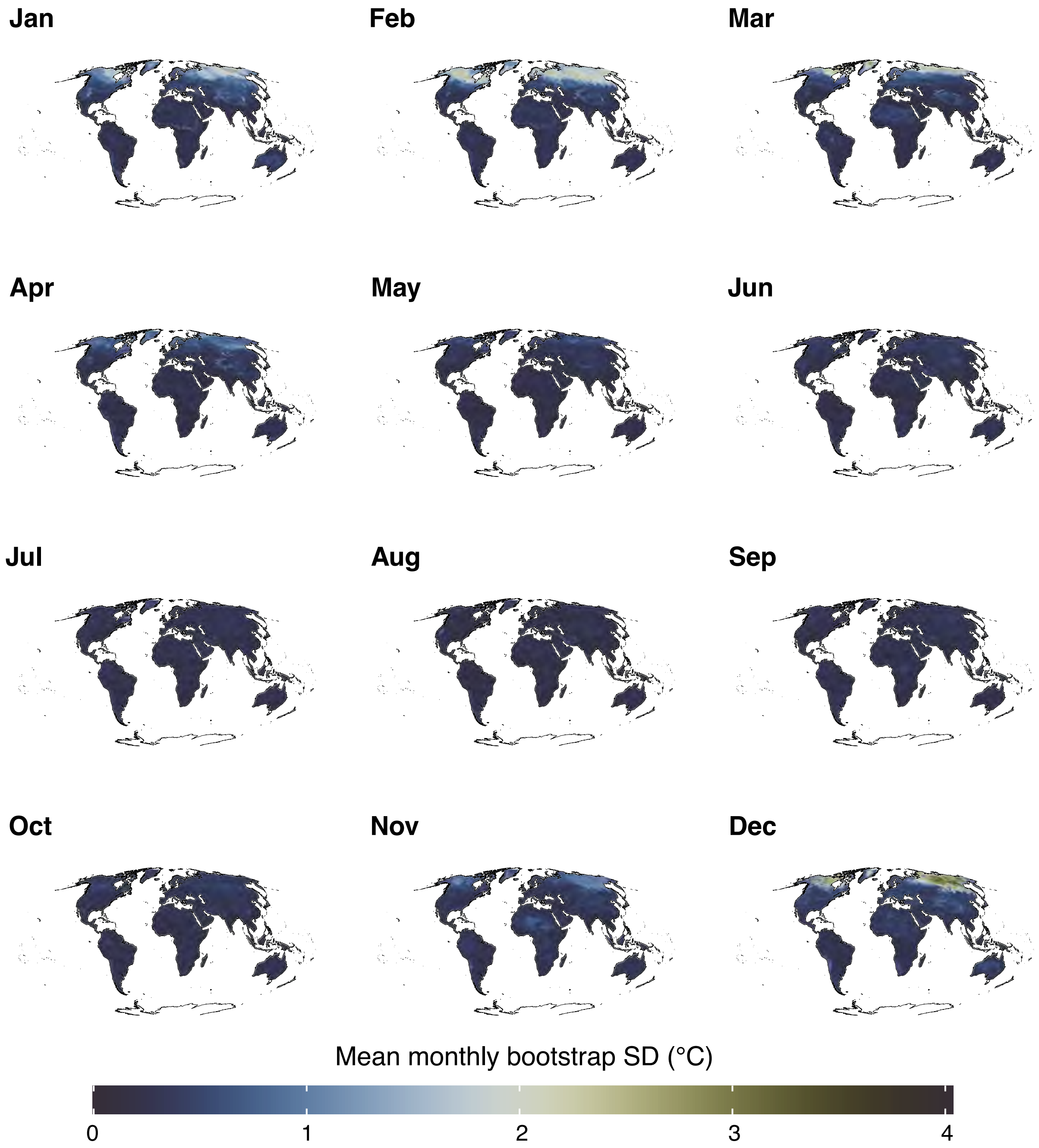


Figure S10: Global monthly maps of bootstrapped standard deviations derived from 100 RF models. Areas with higher values indicate less precise model predictions.

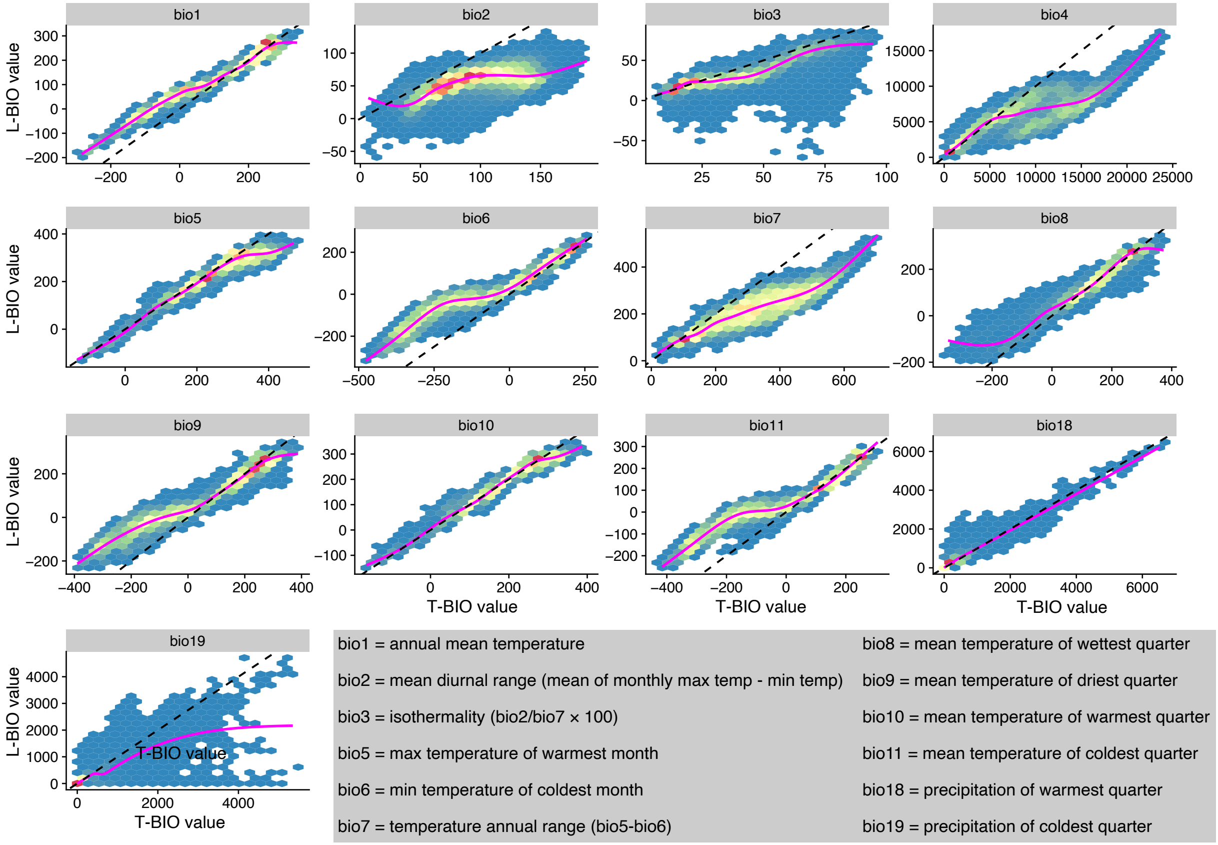


Figure S11: Relationships between original air temperature-based bioclimatic variables (T-BIO) and LSWT-corrected counterparts (L-BIO). Dashed lines show 1:1 relationship, and magenta lines show GAM fits. The definitions of each bioclimatic layer are given in the figure legend. Following common practice, all temperatures are scaled by 100, and precipitation for bio18 and bio19 is in millimeters.

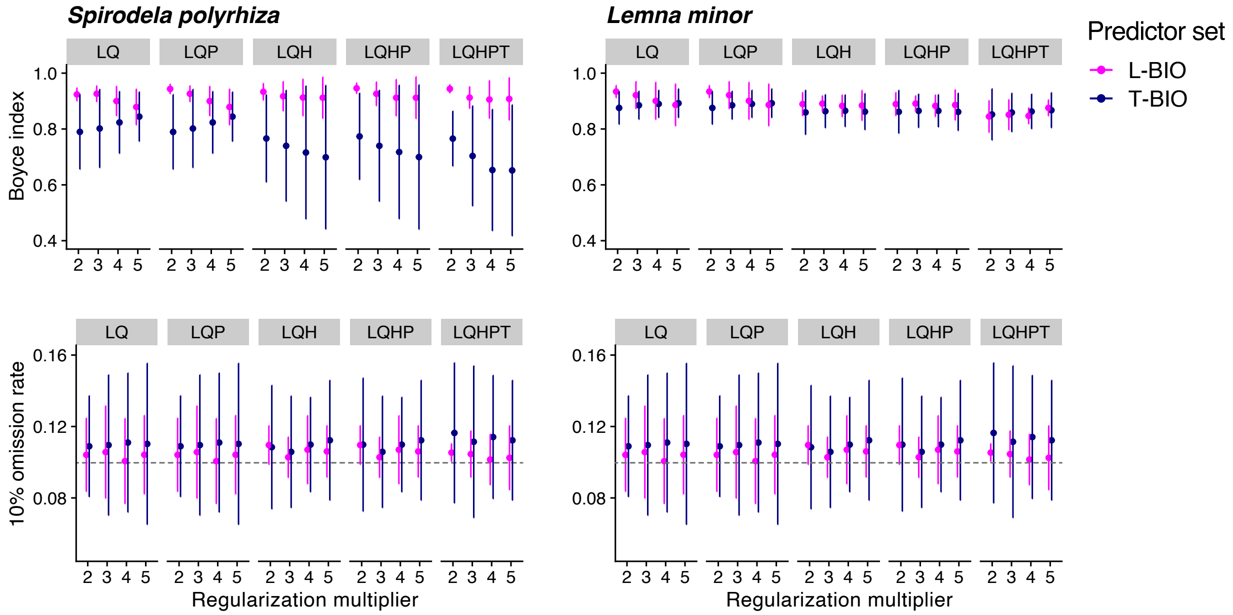


Figure S12: Tuning plots of MaxEnt environmental niche models for the duckweeds *Spirodela polyrhiza* and *Lemna minor*. Each individual model is represented by its own set of regularization multipliers (2-5), feature classes (L = linear, Q = quadratic, H = hinge, P = polynomial, T = threshold), and predictor sets (T-BIO or L-BIO). Upper panels compare Boyce Index values averaged over each model's four CV folds. Values closer to one indicate a better-performing model. Lower panels compare models' 10% omission rates. Values of 0.1 (dashed line) indicate a good model fit.

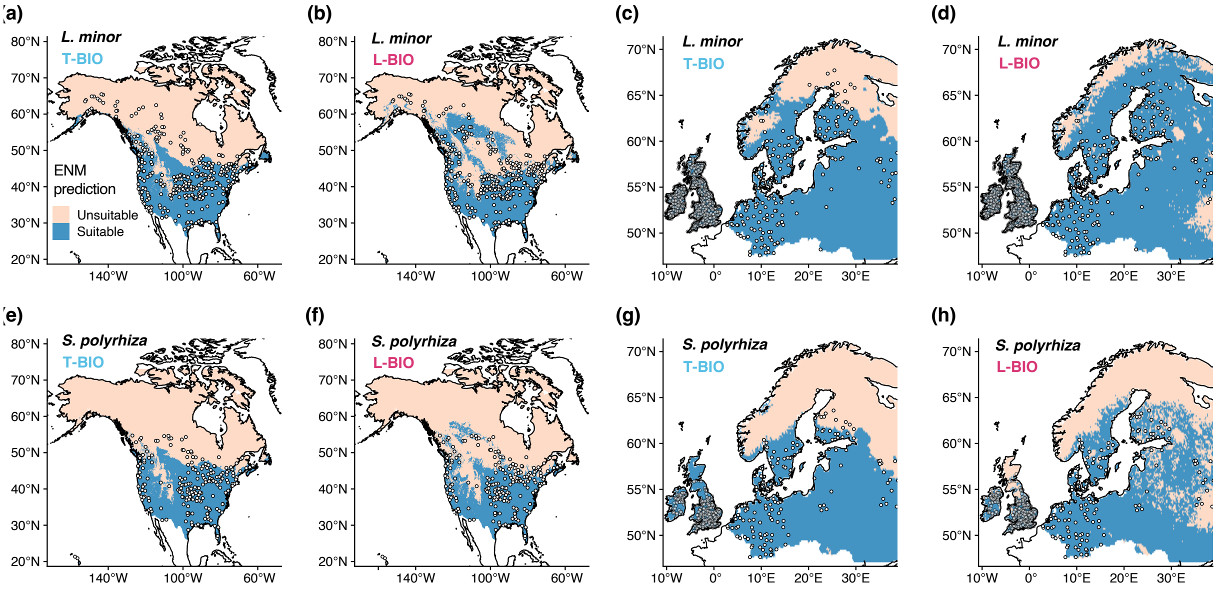


Figure S13: Binary range predictions from best-fit MaxEnt models trained on either the full set of T-BIO or L-BIO predictors. Suitability thresholds were defined as values that omitted the lowest-scoring 10% of observations in cumulative log-log space. While imperfect, note that range limits predicted using L-BIO covariates appear to better reflect the observed margins of observation records for both species.
